# CXCR4-binding PET tracers link monocyte recruitment and endothelial injury in murine atherosclerosis

**DOI:** 10.1101/2020.01.02.892935

**Authors:** Osamu Baba, Andrew Elvington, Martyna Szpakowska, Deborah Sultan, Gyu Seong Heo, Xiaohui Zhang, Hannah Luehmann, Lisa Detering, Andy Chevigné, Yongjian Liu, Gwendalyn J. Randolph

## Abstract

Viral macrophage inflammatory protein 2 (vMIP-II/vCCL2) binds to multiple chemokine receptors, and vMIP-II based PET tracer (^64^Cu-DOTA-vMIP-II: vMIP-II tracer) accumulates at atherosclerotic lesions in mice. The magnitude of ^64^Cu-DOTA-vMIP-II accumulation correlated with monocyte recruitment, as *Apoe^-/-^* mice treated with AAV-mApoE showed PET signal declining as monocyte recruitment subsided. Unexpectedly, monocytes themselves were not the major target of the ^64^Cu-DOTA-vMIP-II tracer. Using fluorescence-tagged vMIP-II tracer, competitive receptor blocking with CXCR4 antagonists, CXCR4-specific tracer ^64^Cu-DOTA-FC131, or CXCR4 staining during disease progression and regression, endothelial cell expression of CXCR4 proved to be the main target of ^64^Cu-DOTA-vMIP-II imaging. Expression of CXCR4 was low in non-plaque areas, but strongly detected on endothelium at the edges of progressing plaques, corresponding to a population of proliferating endothelium and to the location in plaques where monocyte recruitment occurred. Thus, endothelial injury status of plaques is marked by CXCR4 expression and that this injury correlates with the tendency of such plaques to recruit monocytes. Furthermore, our findings suggest PET tracers that, through binding CXCR4, may be useful to monitor plaque injury status.

## Introduction

Atherosclerotic disease is still one of the leading causes of death all over the world even after several medical technologies such as revascularization therapy, including catheter intervention and surgery, and drug treatment have been developed (1). Several imaging tools including ultrasonography, catheter angiography, computed tomography (CT) and magnetic resonance imaging (MRI) have also been developed to detect and assess atherosclerotic plaques (2). However, while many of these tools can detect differences in plaque burden and stenosis of the vascular lumen, measurement of features that predict plaque rupture are not necessarily met by these tools. That is, atherosclerotic plaque rupture, which causes sudden occlusion of an artery and subsequent ischemic tissue injury downstream of the culprit site, does not always occur at severely stenotic lesions (3, 4). Thus, tools that can assess plaque status beyond its overall size are needed (5). In this respect, positron emission tomography (PET) is attractive because of its high sensitivity and its potential for coupling to probes that allow functional evaluation of molecular and cellular aspects. Some specific radioligands are now being used or considered to target the specific pathophysiological process in atherosclerosis (2, 6–8). The identification of imaging tools that predict plaque behavior requires not only application of tools designed to assess specific targets, but follow-up studies to characterize how the tools behave in a complex environment in vivo. Given that the recruitment of monocytes is a signature feature of atherosclerotic plaque activity (9)(10)(11)(12), tools designed to assess monocyte and macrophage activity may be particularly valuable.

Chemokines are a group of about 50 chemotactic heparin-binding cytokines, which are known to be involved in various inflammatory diseases due to their critical roles in directing the movement of circulating leukocytes to sites of inflammation or injury through corresponding chemokine receptors (13). At least 10 chemokine receptors have been identified or implicated in atherosclerotic plaque (14, 15). Viral macrophage inflammatory protein 2 (vMIP-II), also known as vCCL2, is encoded by Kaposi’s sarcoma-associated human herpes virus 8 (HHV-8) and is able to bind to multiple chemokine receptors including CCR1, CCR2, CCR3, CCR5, CCR8, CCR10, XCR1, CX3CR1, CXCR4, and CXCR7 (ACKR3) (16–18)(19), such that it detects collectively the majority of atherosclerosis-related chemokine receptors. Recently, we developed vMIP-II based nanoprobes for PET imaging (^64^Cu-DOTA-vMIP-II), and we documented its uptake in wire injury-induced atherosclerotic plaques (20). However, we did not fully investigate the underlying basis for the signal intensity of the vMIP-II based PET tracer. In particular, it remained unclear which cell populations and chemokine receptors were the major source of the PET signal.

We set out to address that issue herein, with the goal to take into account the possibility that the tracer may not have full access to deeper areas of plaques that might be rich in macrophages and thus might preferentially react with populations available to the blood circulation, such as monocytes. To that end, we designed a protocol that took advantage of the fact that monocyte recruitment to plaques in *Apoe^-/-^* mice is readily modulated by restoring apoE expression using viral vectors (12). Indeed, monocyte recruitment is dramatically suppressed by apoE expression in *Apoe^-/-^* mice before macrophage burden decreases. Thus, use of this approach that kinetically separates changes in monocyte recruitment with overall macrophage burden would allow us to ask whether the broad spectrum vMIP-II based PET tracer could be used to read out one or the other of these parameters.

Here, we demonstrate that the PET signal derived from the ^64^Cu-DOTA-vMIP-II tracer indeed correlates with monocyte recruitment into atherosclerotic plaque rather than macrophage burden. However, in the process of exploring mechanism of action, we uncover the surprising result that ^64^Cu-DOTA-vMIP-II signal in *Apoe^-/-^* mice primarily targeted CXCR4 on endothelium. This finding then opened the door for further experiments that revealed that CXCR4 expression is induced on the endothelium at the margins of plaques where endothelial permeability and proliferation is highest. Through the studies presented herein, we argue that CXCR4 serves as an important marker of injury in the endothelium and that this status is closely linked to monocyte recruitment. Considering that CXCR4 locus in human has been reported to be as a key determinant in coronary heart disease (21) these new findings underscore the possibility that CXCR4-targeted PET tracers or perhaps the ^64^Cu-DOTA-vMIP-II tracer may report critical information about plaque status in a clinical setting.

## Results

### ^64^Cu-DOTA-vMIP-II signal correlates with monocyte recruitment rather than plaque size or macrophage burden in atherosclerotic plaques

We first confirmed that ^64^Cu-DOTA-vMIP-II tracer (vMIP-II tracer) accumulates at atherosclerotic plaques by comparing uptake of the tracer assessed by autoradiography with plaque distribution assessed by ORO staining in whole thoracic aortas from *Apoe^-/-^* mice fed high fat diet (HFD) for 10 weeks (Fig. 1A). As shown in Fig. 1A, the high tracer localization was observed at the arch region where high lipid deposition was determined, indicating the binding of tracer to plaque. Next, we used a plaque regression model to compare the uptake of ^64^Cu-DOTA-vMIP-II tracer and several biological parameters related to atherosclerosis side by side at several time points (12). We treated *Apoe^-/-^* mice on HFD with AAV8 vector encoding murine *Apoe* (AAV-mApoE vector) to complement *Apoe* expression or PBS as control. Then, PET imaging was carried out at baseline, and 2 or 4 weeks after transduction. Aortas were subsequently removed to measure tracer biodistribution by scintillation counting (Fig. S1A). In this model, plasma total cholesterol level decreased and reached nadir at ∼200 mg/dL 1 week after AAV-mApoE transduction (Fig. S1B). There was no difference in atherosclerotic plaque area in this time period (Fig. 1B-C), but it was significantly decreased 8 weeks after treatment (Fig. S1C). Monocyte recruitment was assessed by labeling Ly6C^hi^ monocytes with fluorescent polystyrene particles (Fig. S1D-E) (12). Counting of bead-labeled cells in plaques confirmed reduction of monocyte recruitment by 2 weeks after transduction with AAV-mApoE and time points beyond (Fig. 1D). Macrophage accumulation analyzed by CD68 staining did not change at 2 weeks but decreased at 4 weeks post treatment (Fig. 1E-F). MOMA-2 staining showed the consistent result with CD68 staining (Fig. S1F). Therefore, in this model, plasma total cholesterol is decreased at first, then monocyte recruitment, macrophage accumulation and plaque size are decreased in turn. Uptake of ^64^Cu-DOTA-vMIP-II tracer at the aortic arch, assessed by PET imaging, was decreased by 2 weeks of AAV-mApoE treatment (Fig. 1G-H). Biodistribution in aortas was likewise decreased (Fig. S1G). Linear regression analysis also showed the positive correlation between PET signal and monocyte recruitment (Fig. S1H). Thus, changes in monocyte recruitment but not changes in plaque macrophage burden or plaque area, tracked most closely with the aortic uptake of ^64^Cu-DOTA-vMIP-II tracer.

**Fig.1.**
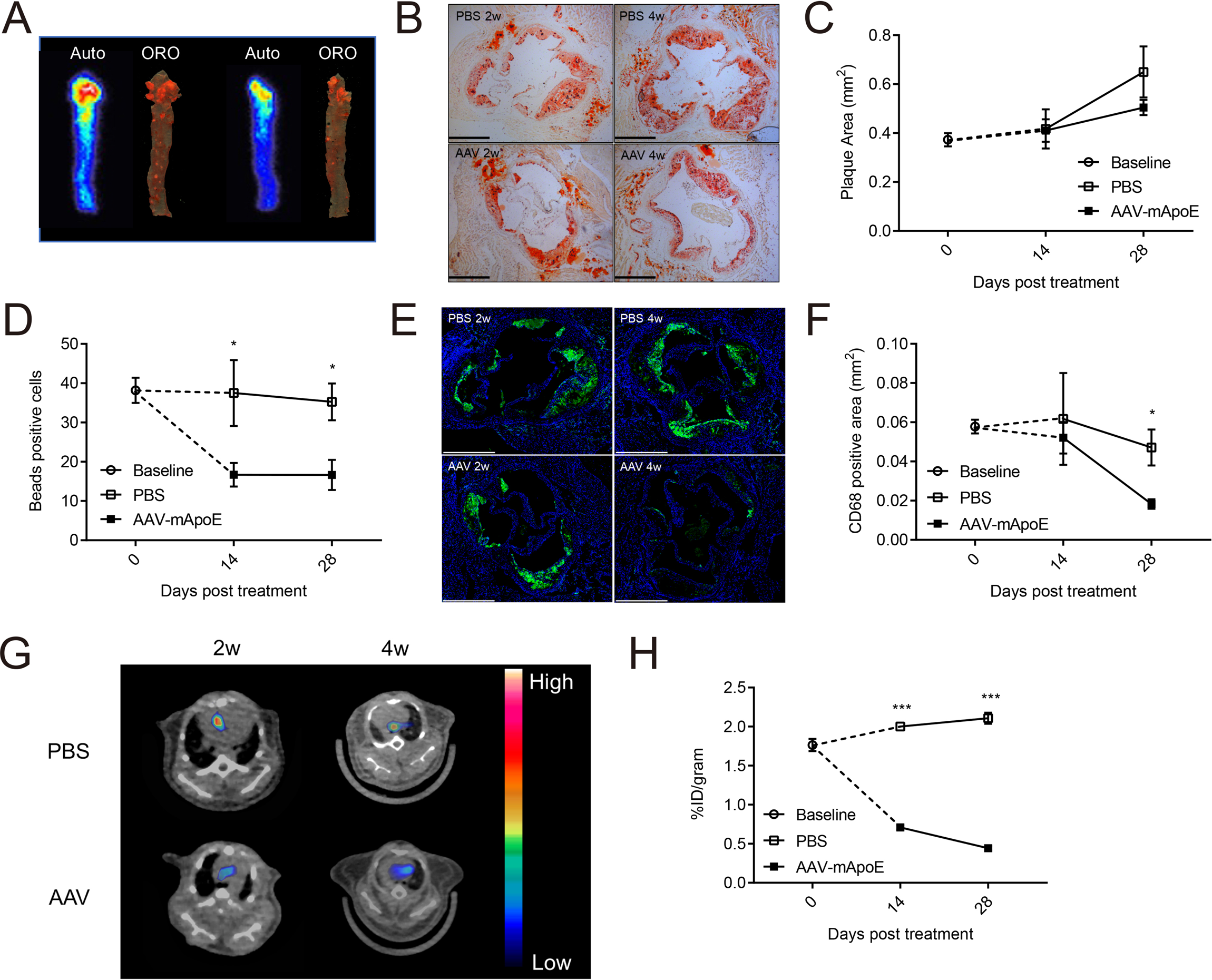
PET signal derived from ^64^Cu-DOTA-vMIP-II tracer correlates with monocyte recruitment into atherosclerotic plaques. **(A)** Autoradiography of ^64^Cu-DOTA-vMIP-II (left: indicated as auto) and corresponding oil red O staining (right: indicated as ORO) of whole thoracic aortas. Time course of **(B-C)** plaque areas in aortic sinuses, **(D)** beads positive cells in plaques **(E-F)** CD68 positive areas, and. **(G)** Representative ^64^Cu-DOTA-vMIP-II PET images at aortic arches and **(H)** quantification of PET signal (n=4-8 per point). All data are shown as means ± SEM. *p<0.05, ***p<0.001 by Student’s *t* test.

### In vivo distribution of ^64^Cu-DOTA-vMIP-II tracer

To further assess the distribution of ^64^Cu-DOTA-vMIP-II tracers *in vivo*, we made a Cy5-conjugated non-radioactive vMIP-II tracer (Cy5-vMIP-II). Within 1 h after injection, this tracer was found in blood of HFD-fed *Apoe^-/-^* mice mainly bound to monocytes (Ly6C^hi^>Ly6C^lo^) and neutrophils, with lesser binding to T and B cells in a dose-dependent manner (Fig. 2A). Here, most of Cy5-conjugated tracers lined the plaque surface and colocalized with CD31^+^ endothelial cells (ECs), rather than monocytes/macrophages, in the plaque sections (Fig. 2B). Flow cytometric analysis of atherosclerosis-bearing aortas confirmed binding to ECs and recruited monocytes and neutrophils, but not macrophages (Fig. 2C, S2A). The tracer may have limited access to plaque macrophages, as it may not deeply penetrate into plaques. Another dye-conjugated vMIP-II tracer (CF640R-vMIP-II) also showed its binding capacity for peripheral monocytes and neutrophils (Fig S2B), and accumulation at atherosclerotic plaques in whole mount imaging of the thoracic aorta (Fig. S2C) and in aortic ECs (Figure S2D). We conclude that there are two likely sources of PET signal arising from the use of vMIP-II tracer in atherosclerotic mice: (1) tracer-bound blood leukocytes, neutrophils or monocytes, recruited into atherosclerotic lesions from blood and (2) direct binding to plaque ECs.

**Fig.2.**
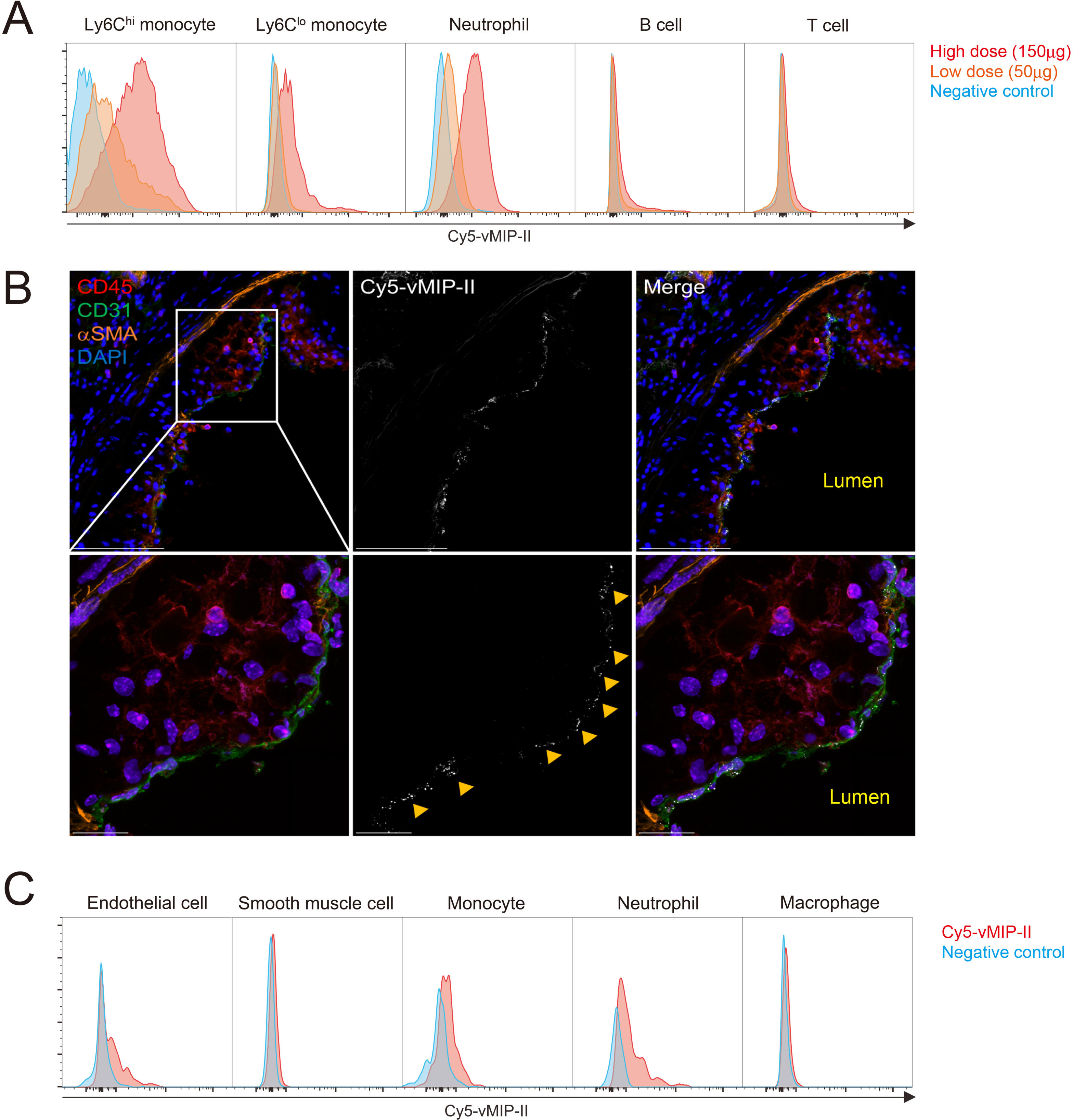
Cy5-conjugated vMIP-II tracer binds to blood monocytes and neutrophils and to plaque endothelial cells (ECs) *in vivo*. **(A)** Histogram of Cy5^+^ leukocyte populations in peripheral blood of *Apoe^-/-^* mice 1 h after injection of high dose (150 μg) or low dose (50 μg) of Cy5-vMIP-II tracer. Negative control is each cell population from *Apoe^-/-^* mice injected PBS. **(B)** Distribution of Cy5-vMIP-II tracer (150 μg/mouse) in atherosclerotic plaques examined at low (upper) and high (middle) magnification. Accumulation of the tracer is indicated by yellow arrows. Scale bar: 100 μm (upper) and 20 μm (bottom) **(C)** Histogram of Cy5^+^ cell populations in the aorta of *Apoe^-/-^* mice 1 h after injection of 150 μg of Cy5-vMIP-II tracer. Negative control is each cell population from *Apoe^-/-^* mice injected PBS.

### Recruitment of tracer-bound neutrophils and monocytes are not major source of PET signal of ^64^Cu-DOTA-vMIP-II tracer

To assess the concept that the tracer directly detected monocyte or neutrophil recruitment to plaque, we depleted peripheral monocytes and neutrophils using depleting antibodies in *Apoe^-/-^* mice fed HFD. Neutrophils were selectively depleted by use of anti-Ly6G mAb, and both neutrophils and Ly6C^hi^ monocytes were depleted by use of Gr-1 mAb recognizing Ly6C and Ly6G (Fig. S3A-B). Ly6C^lo^ monocytes were not depleted in this experiment. However, more than half of the monocytes recruited to atherosclerotic plaques are Ly6C^hi^ monocytes (22, 23), and the preferential binding of dye conjugated-vMIP-II to Ly6C^hi^ monocytes predisposes to detecting this subset (Fig. 2A, S2B). Only a mild reduction of the uptake was observed 24 h after treatment even after both neutrophil and Ly6C^hi^ monocyte were depleted, although it is statistically significant (Fig. 3A-B). Biodistribution of the tracer in aortas was also mildly decreased after monocyte and neutrophil depletion (Fig. S3C). Therefore, direct recognition of monocytes or neutrophils can explain only a minority of the reduction of reactivity of vMIP-II tracer in regressing plaques.

### Neither change of glycosaminoglycan content in atherosclerosis nor nonspecific tracer recruitment into plaques accounts for tracer reactivity

Several chemokines including vMIP-II can also bind to glycosaminoglycans (GAGs) (24, 25) which are known to have important roles in leukocyte adhesion and recruitment, and vessel permeability. Moreover, the amount of each type of GAGs changes respectively in atherosclerotic artery (26–28). Thus, we considered the possibility that plaque GAGs that might bind to the ^64^Cu-DOTA-vMIP-II tracer were increased in atherosclerotic plaques actively recruiting monocytes. To this end, we measured sulfated GAG content in aortas from *Apoe^-/-^* mice with or without AAV treatment, but we found no reduction in total GAG content 2 weeks after AAV-mApoE treatment (Fig. 3C). Moreover, the content of subtypes of GAGs (O- and N-sulfated GAGs) also did not change (Fig. S3D).

**Fig.3.**
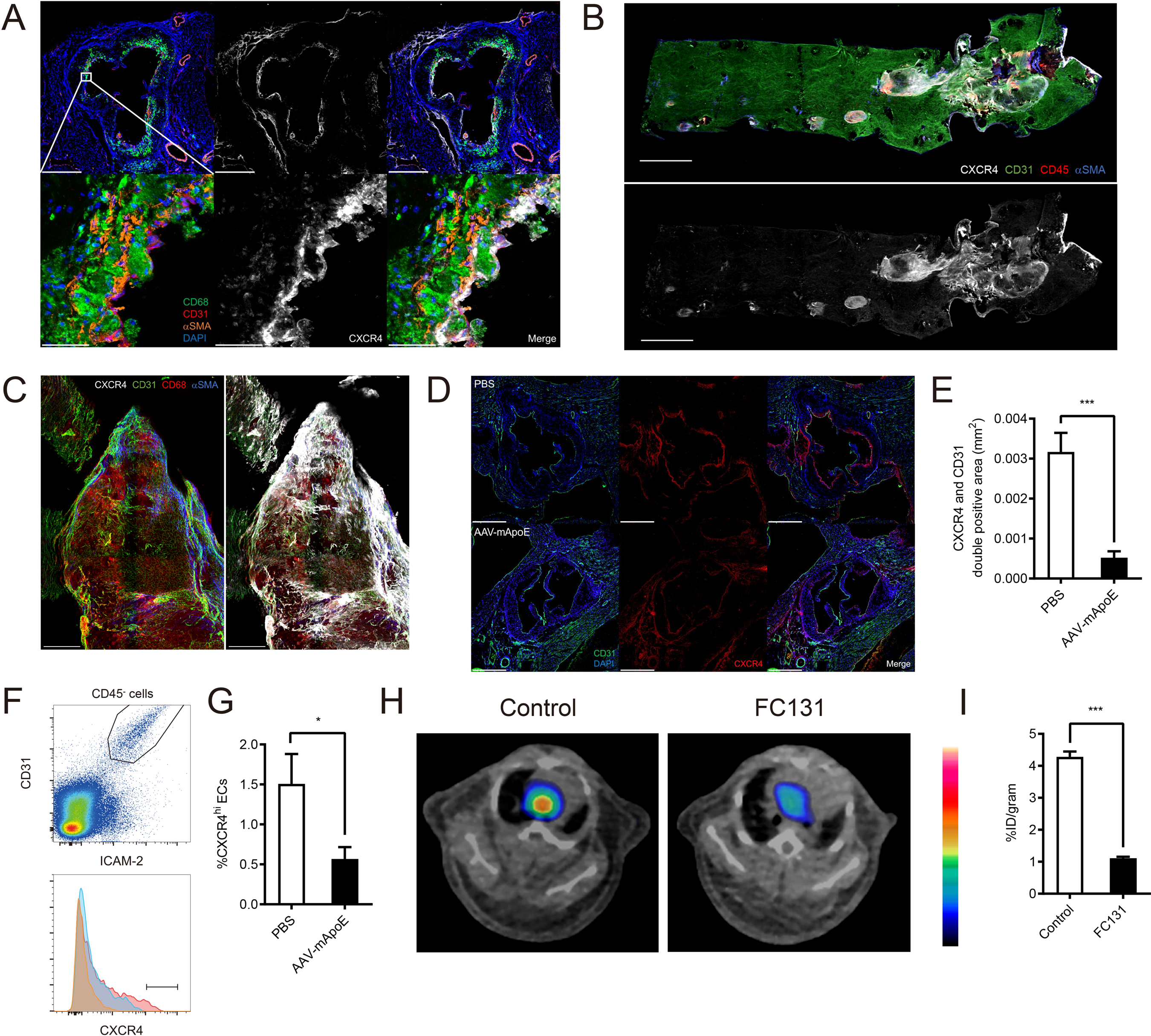
Recruitment of tracer-bound leukocytes, non-specific binding of the tracer, or change in plaque glycosaminoglycan content do not account for the reduction of PET signal in regressing plaques. **(A)** Representative ^64^Cu-DOTA-vMIP-II PET images and **(B)** quantification (n=3-4 per group) at aortic arches of *Apoe^-/-^* mice on HFD with or without leukocyte depletion. *p<0.05 by 1-way ANOVA with the Turkey post hoc test. **(C)** Sulfated glycosaminoglycan (sGAG) content in whole thoracic aortas from *Apoe^-/-^* mice with or without 2 weeks of AAV-mApoE treatment (n=4-5 per group). **(D)** Biodistribution of ^64^Cu-DOTA-PEG in aortas from *Apoe^-/-^ mice* with or without 2 weeks of AAV-mApoE treatment, or WT mice (n=4-5 per group). All data are shown as means ± SEM.

We also considered the possibility that the tracer identity is unimportant and that the greater intensity in progressing plaques reflects higher endothelial permeability to nonspecific substances (29, 30). To assess this, we employed a tracer which had the similar molecular weight (∼10 KDa) as the vMIP-II tracer but without the chemokine receptor binding activity (^64^Cu-DOTA-PEG). We then measured biodistribution of this tracer in aortas from *Apoe^-/-^* mice with or without AAV treatment, or WT mice as control. Although tracer accumulation in aortas tended to increase in *Apoe^-/-^* mice compared with WT mice, there was no difference between *Apoe*^-/-^ mice with and without AAV-mApoE treatment (Fig. 3D). Moreover, the uptake of nonspecific ^64^Cu-DOTA-PEG tracer was negligible compared with ^64^Cu-DOTA-vMIP-II tracer. We thus conclude that ^64^Cu-DOTA-vMIP-II tracer uptake in atherosclerotic plaque was mainly due to the binding to chemokine receptors overexpressed on aortic endothelial cells.

### ^64^Cu-DOTA-vMIP-II tracer mainly reports CXCR4 expression on plaque ECs that varies <u>with disease activity</u>

Some vascular ECs highly express CXCR4 (31, 32), which in turn has binding capacity for vMIP-II (16–18). Given that we visualized tracer binding to endothelium (Fig. 2B, S2D), it seemed possible that vMIP-II tracer binds to plaque ECs through CXCR4. Staining of aortic sinus from *Apoe^-/-^* mice on HFD showed that CXCR4 was expressed in not only ECs but also macrophages and smooth muscle cells. However, interestingly, only cells located close to the surface of the plaque expressed CXCR4 (Fig. 4A). We also assessed distribution of CXCR4 in thoracic aortas using whole-mount staining, which revealed that CXCR4 was mainly expressed at atherosclerotic lesions, especially at the edge of plaques (Fig. 4B-C) where monocytes are known to be mainly recruited (33). Moreover, CXCR4^+^ CD31^+^ areas were significantly reduced in regressing plaques (Fig. 4D-E). Flow cytometric analysis also showed a reduction in CXCR4^hi^ ECs 2 weeks post AAV-mApoE treatment (Fig. 4F-G). The recent literature demonstrated that vMIP-II also has a binding capacity for one of the atypical chemokine receptors CXCR7 (ACKR3) (19) which is expressed in ECs (34). However, CXCR7 expression on aortic ECs did not change post AAV-mApoE treatment (Fig. S4C-D). Most compellingly, blocking of CXCR4 by CXCR4-selective antagonist FC131 (35), which uses the same binding site with vMIP-II (36), dramatically reduced PET signal of vMIP-II tracer at aortic arches of *Apoe^-/-^* mice on HFD (Figure 4H-I), as also observed when analyzed by biodistribution (Fig. S4E), indicating that CXCR4 is a major, accessible target for vMIP-II tracer. We also demonstrated that our fluorescence-conjugated versions of vMIP-II bound to both human CXCR4 and CXCR7 expressing cells *in vitro* but displayed a higher affinity for CXCR4 (Fig. S4A-B).

**Fig.4.**
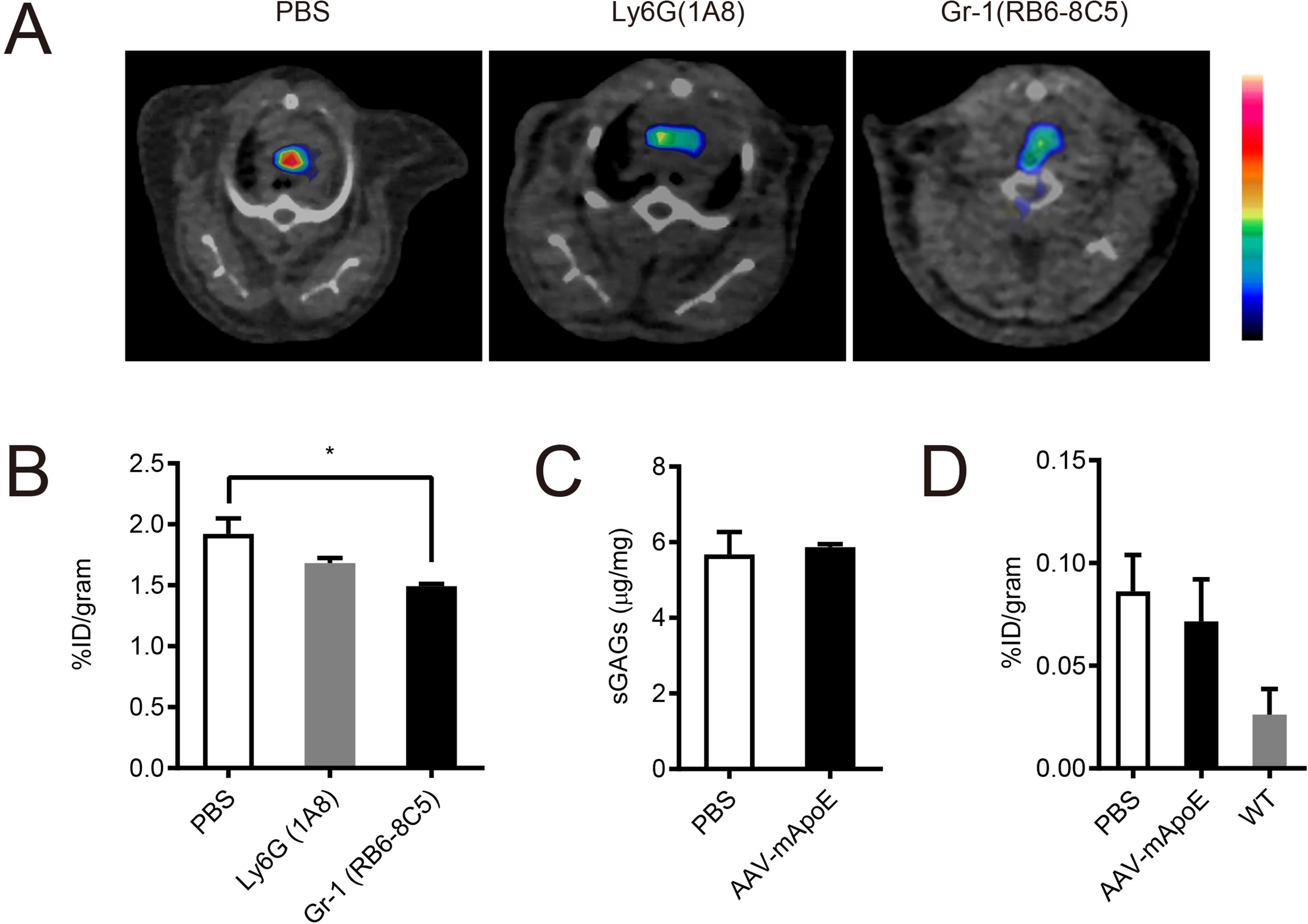
CXCR4 accounts substantially for the PET signal arising from the ^64^Cu-DOTA-vMIP-II tracer. **(A)** CXCR4 expression in aortic sinus from *Apoe^-/-^* mice on HFD. Scale bar: 500 μm (upper) and 50 μm (bottom) **(B)** CXCR4 distribution in whole thoracic aorta from *Apoe^-/-^* mice on HFD. Scale bar: 1 mm **(C)** CXCR4 distribution in the plaque at the aortic arch from *Apoe^-/-^* mice on HFD. Scale bar: 200 μm **(D)** Representative images of CXCR4 expression in aortic sinus from *Apoe^-/-^* mice on HFD after 2 weeks of PBS (upper) or AAV-mApoE (bottom) treatment. Scale bar: 500 μm, and **(E)** quantification of CXCR4^+^ CD31^+^ areas (n=6 per group). **(F)** Representative histogram of CXCR4 expression on aortic ECs from *Apoe^-/-^* mice on HFD after 2 weeks of PBS (red) or AAV-mApoE (blue) treatment. Yellow histogram is isotype control. **(G)** The percentage of CXCR4^hi^ ECs in total aortic ECs with or without 2 weeks of AAV-mApoE treatment (n=7 per group). **(H)** Representative ^64^Cu-DOTA-vMIP-II PET images and **(I)** quantification (n=5 per group) at aortic arches of *Apoe^-/-^* mice on HFD with or without CXCR4 blocking by 0.5 mg of FC131 (n=5 per group). All data are shown as means ± SEM. *p<0.05, ***p<0.001 by Student’s *t* test.

### CXCR4 expression in atherosclerotic lesions marks sites where endothelial permeability is prominently enhanced

Moreover, vascular permeability is known to be increased at atherosclerotic lesions (29, 30). Endothelial CXCR4 attenuates vascular permeability through enhancement of VE-cadherin expression and stabilization of junctional VE-cadherin complexes (21). Therefore, we wondered whether endothelial CXCR4 expression is upregulated to compensate for enhanced vascular permeability at atherosclerotic lesions. To examine this hypothesis, we first assessed distribution of permeability in thoracic aorta lesions *of Apoe^-/-^* mice on HFD by *in vivo* injection of Evans Blue dye. As expected, Evans Blue leakage mainly occurred at atherosclerotic lesions but interestingly especially at the edge of plaques (Fig. 5A), with dye distribution occurring in similar areas as CXCR4 distribution (Fig. 4C). Indeed, co-staining of CXCR4 demonstrated Evans Blue leakage around CXCR4-expressing endothelial areas (Fig. 5B). Since CXCR4 blocking or endothelial specific CXCR4 deficiency was earlier reported to enhance Evans Blue leakage from aortic arches (21), this colocalization likely reflects a compensatory attempt of the endothelium to combat elevated permeability.

**Fig.5.**
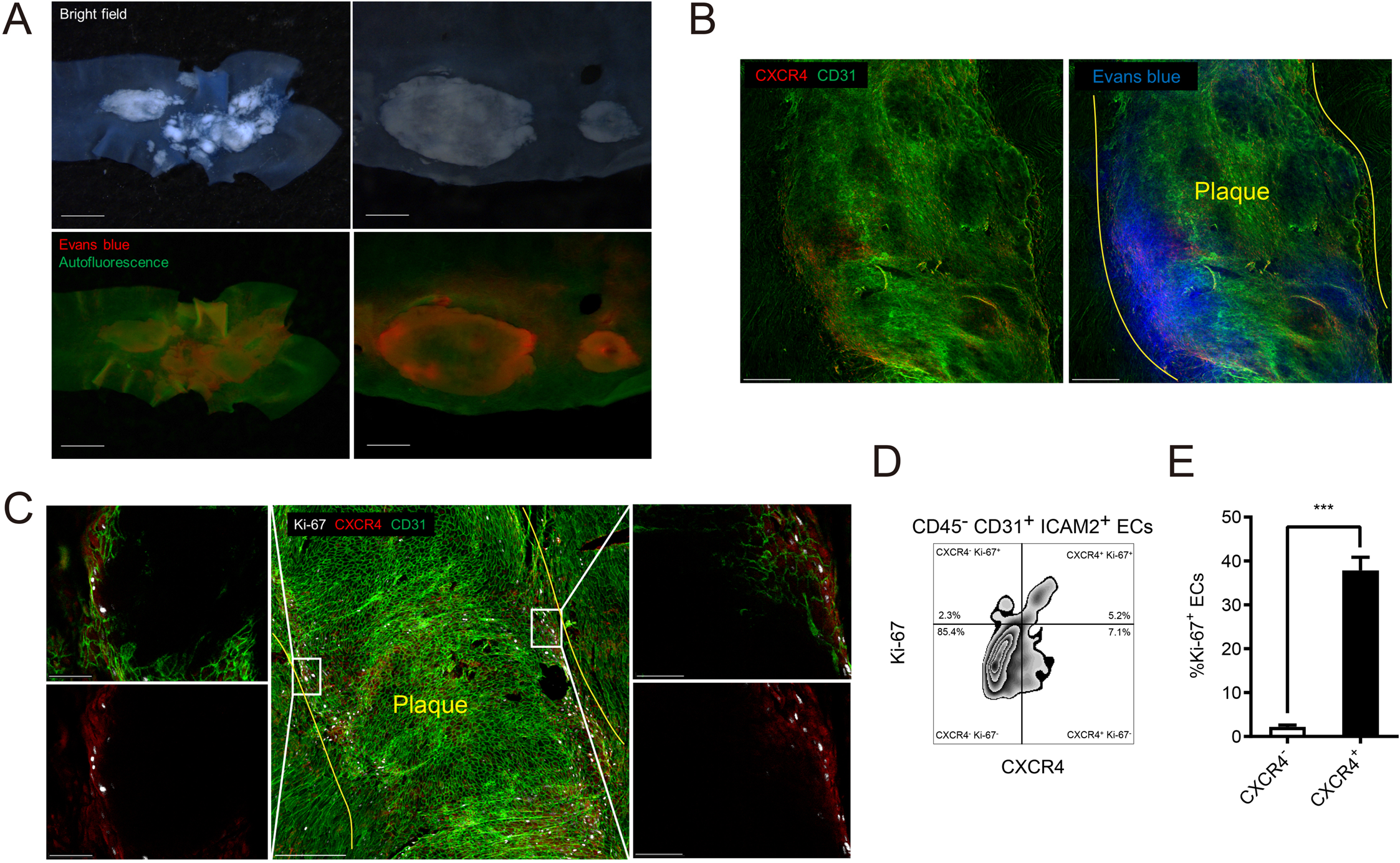
Vascular permeability is increased along the margins of atherosclerotic plaques where CXCR4-expressing ECs are proliferating. **(A)** Bright field (upper) and fluorescent (bottom) stereoscope images of an aortic arch (left) and plaques in a descending aorta (right) from *Apoe^-/-^* mice injected 200 μL of 0.5% Evans blue. Scale bar: 1 mm (left) and 500 μm (right) **(B)** CXCR4 distribution in a plaque of *Apoe^-/-^* mice injected evans blue. Yellow lines indicate the edged of the plaque. Scale bar: 200 μm **(C)** Ki-67 and CXCR4 distribution in atherosclerotic plaques at an aortic arch with low (middle) and high (left and right) magnification. Yellow lines indicate the edged of the plaque. Scale bar: 200 μm (middle) and 50 μm (left and right). **(D)** The scheme for gating of Ki-67 or CXCR4 positive ECs. **(E)** The percentage of Ki-67 positive cells in CXCR4 positive or negative ECs (n=6 per group). Data are shown as means ± SEM. ***p<0.001 by paired *t* test.

### Analysis of endothelial changes in proliferation and turnover

_Endothelial denudation and turnover are promoted at atherosclerotic lesions (37, 38). After vascular injury, CXCR4 enhances endothelial proliferation and re-endothelialization while Ki-67^+^ ECs are reduced in endothelial-specific CXCR4-deficient mice after endothelial injury (39). Therefore, we hypothesized that endothelial CXCR4 may promote recovery from endothelial injury or denudation especially at the edge of plaques by promoting proliferation. To assess this possibility, we stained Ki-67 as a proliferation marker in whole thoracic aorta. Ki-67^+^ cells were mainly located at the edge of atherosclerotic plaques and were CXCR4^+^ ECs (Fig. 5C). Flow cytometric analysis confirmed that CXCR4^+^ aortic ECs were more proliferative (Fig. 5D-E). These data are consistent with the previously reported protective role that CXCR4 has in endothelial repair and in combatting heightened permeability (39). Despite the potentially direct protective effects of CXCR4, ECs that have induced CXCR4 would be those facing adverse conditions that lead to heightened permeability and need for repair in the first place and thus CXCR4^+^ ECs may also express pro-inflammatory markers. Indeed, CXCR4^+^ HUVEC upregulate MCP-1 (CCL2) (40), a chemokine that can directly induce monocyte recruitment. Consistent with this, MCP1 expression was increased in CXCR4 expressing aortic ECs from *Apoe^-/-^* mice on HFD (Fig. S5A-B). Thus, our data, together with existing literature, would suggest that CXCR4 expression on ECs correlates with endothelial loss of homeostasis and thus appears to mark ECs that actively promote monocyte recruitment, while also having intrinsic protective roles to restore homeostasis.

### CXCR4 specific PET tracer also accumulates in plaque ECs and shows a reduction of PET signal in regressing plaques

To further investigate the hypothesis that CXCR4 signal reports the level of endothelial injury in atherosclerosis, we compared ^64^Cu-DOTA-vMIP-II to ^64^Cu-DOTA-FC131, a CXCR4 specific tracer. Although PET signals arising from CXCR4-specific tracer is already reported in atherosclerosis (41–44), the plaque characteristics, or biological parameters of plaque activity, that may or may not correlate with its PET signal are still not fully understood. Similar to studies with the vMIP-II tracer (Fig. 1), attenuation of CXCR4-specific PET signal and biodistribution in thoracic aortas were observed 2 weeks post AAV-mApoE treatment (Fig. 6A-B, Fig. S5C), when monocyte recruitment was reduced by plaque area and macrophage burden had not changed. We also detected accumulation of the tracer in other organs, as expected, where the apoE vector reduced binding only in the aorta for both the CXCR4-specific tracer and the vMIP II tracer (Fig. 6A., Fig. S5C). Dye-conjugated CXCR4 specific tracer (CF640R-FC131) also accumulated in plaque ECs (Fig. 6C) and bound to peripheral monocytes and less to neutrophils and B cells (Figure S4D), which is comparable with CXCR4 expression in peripheral leukocytes (Figure S4E). These data indicate that CXCR4-specific tracer can be a useful tool to detect endothelial status in plaques, including sufficient levels of injury to promote monocyte recruitment.

**Fig.6.**
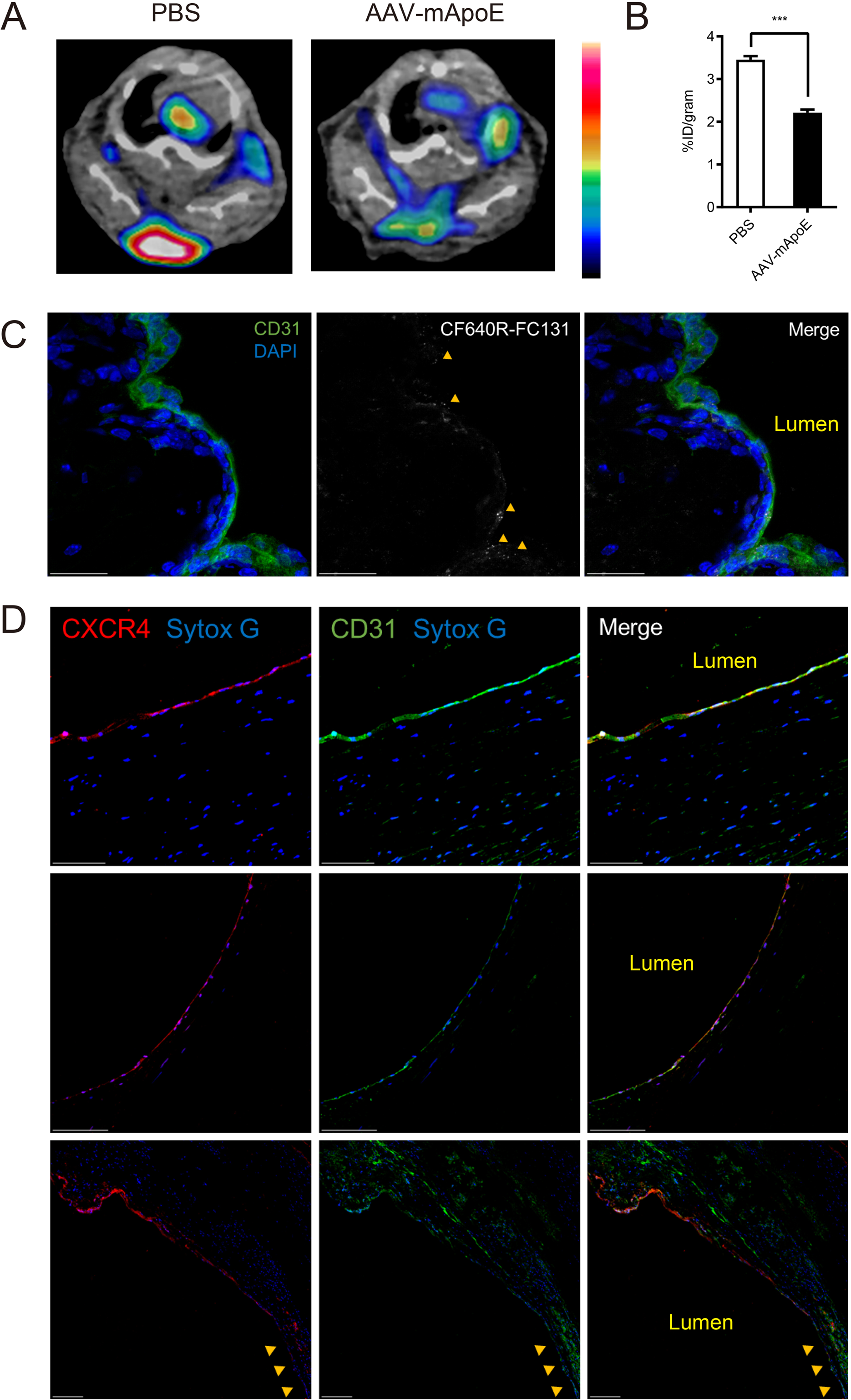
FC131-based CXCR4-specific tracer mirrors the behavior of vMIP-II tracer in regressing versus progressing atherosclerotic plaques. **(A)** Representative ^64^Cu-DOTA-FC131 images at aortic arches of *Apoe^-/-^* mice on HFD 2 weeks post PBS or AAV-mApoE treatment and **(B)** quantification of PET signal (n=6 per group). Data are shown as means ± SEM. **p<0.01 by Student’s *t* test. **(C)** Accumulation of CF640R-FC131 tracer (40 μg/mouse) in an atherosclerotic plaque. Accumulation of the tracer is indicated by yellow arrows. Scale bar: 20 μm **(D)** CXCR4 expression in the human plaque from 2 different specimens stemming from peripheral arterectomies. Two specimens are from one subject’s femoral artery (upper and bottom) and another from an anterior tibialis artery (middle). Non-plaque area in the artery lumen is indicated by yellow arrows (bottom). Scale bar: 100 μm (upper and middle), 200 μm (bottom image).

### CXCR4 is also expressed in human plaque ECs

CXCR4 staining in human atherosclerotic plaques revealed its expression on the surface of the plaque, including colocalization with CD31^+^ ECs (Fig. 6D). Moreover, consistent with our mouse data, its expression was observed at the plaque shoulder and it was low at the adjacent non-plaque area (Fig. 6D). We also confirmed this colocalization using different clone of CXCR4 mAb (Fig. S5F).

## Discussion

We had earlier developed the vMIP-II based PET tracer ^64^Cu-DOTA-vMIP-II, which accumulates at atherosclerotic lesions in mice (20). Based on the nature of vMIP-II, this tracer can bind to a series of chemokine receptors, including most atherosclerosis-related chemokine receptors (16–18). Initially, it seemed reasonable to expect that this tracer might react broadly with plaque macrophages in mouse models of atherosclerosis. However, we were also cognizant of the possibility that some regions of plaque might be inaccessible to the tracer and thus we set out to determine if the tracer might, in fact, be most accessible to monocytes and thereby serve as a beacon of the relative tendency of monocytes to enter atherosclerotic plaques. Indeed, by employing a model system in which monocyte recruitment can be modulated before changes in plaque burden occur, we were able to determine that vMIP-II tracer accumulation in the artery wall of mice with experimental atherosclerosis is a reliable correlate to monocyte recruitment, which in turn is a strong correlate to disease activity.

We were surprised, however, that CXCR4 expressed on plaque ECs was the main target of vMIP-II tracer. Many of the chemokine receptors that would be targets of the vMIP-II tracer are expressed by monocytes and macrophages, including CXCR4 observed earlier on plaque macrophages. Yet, our data suggest that macrophages in plaques are rather inaccessible to our chemokine receptor targeting tracers. We argue that this problem is due to inadequate access of the tracer to the deeper regions of the plaque, whereas the endothelial is quite accessible and thus dominates in signal. Monocytes in the blood are also accessible to the tracer, but the pace of recruitment of monocytes, even during higher levels of recruitment, may also be too low to compete with endothelial cell binding in overall magnitude.

When our data unexpectedly pointed to the plaque endothelium as a major target, we began to consider CXCR4 as the dominant ligand for the vMIP-II tracer. Indeed, we were able to block vMIP-II tracer signal with CXCR4-specific antagonist. CXCR4 is known to be expressed in ECs (31, 32), albeit not expressed under all conditions. We did not directly demonstrate how endothelial CXCR4 is associated with monocyte recruitment in atherosclerosis. However, based on the literature that re-endothelialization of arteries, endothelial proliferation (39), and reduced atherosclerosis (21) is linked to CXCR4 expression and function, we argue that CXCR4 itself has a protective role. Additional recent RNA sequencing data analysis of locally regenerating ECs to recover from endothelial injury *in vivo* shows enhancement of CXCR4 expression (45). Endothelial CXCR4 is also reported to be athero-protective through attenuating vascular permeability by enhancing VE-cadherin expression and stabilizing VE-cadherin complexes (21). However, despite its likely protective functional role, the induction of its expression is known to be a consequence of endothelial injury (39) and thus its expression correlates with active plaque. That our study shows that it is most highly expressed at the margins of plaque, where it is known that monocyte recruitment is most dominant (33) and where we demonstrate there is the most endothelial cell proliferation and permeability fits with the concept that its expression signifies endothelial response to injury. We also found that CXCR4 expressing aortic ECs are more likely to express MCP-1. This expression of MCP-1 does not mean that endothelial CXCR4 signaling per se is athero-promoting, but rather that its expression is associated with a loss of homeostasis that is associated with inflammation, even though the net effect of CXCR4 signaling per se is athero-protective (21).

Overall, we propose that CXCR4 is upregulated in plaque ECs at the edge or shoulder of the plaque to compensate increased vascular permeability through promotion of re-endothelialization and probably VE-cadherin pathways (Fig. 7).

**Fig.7.**
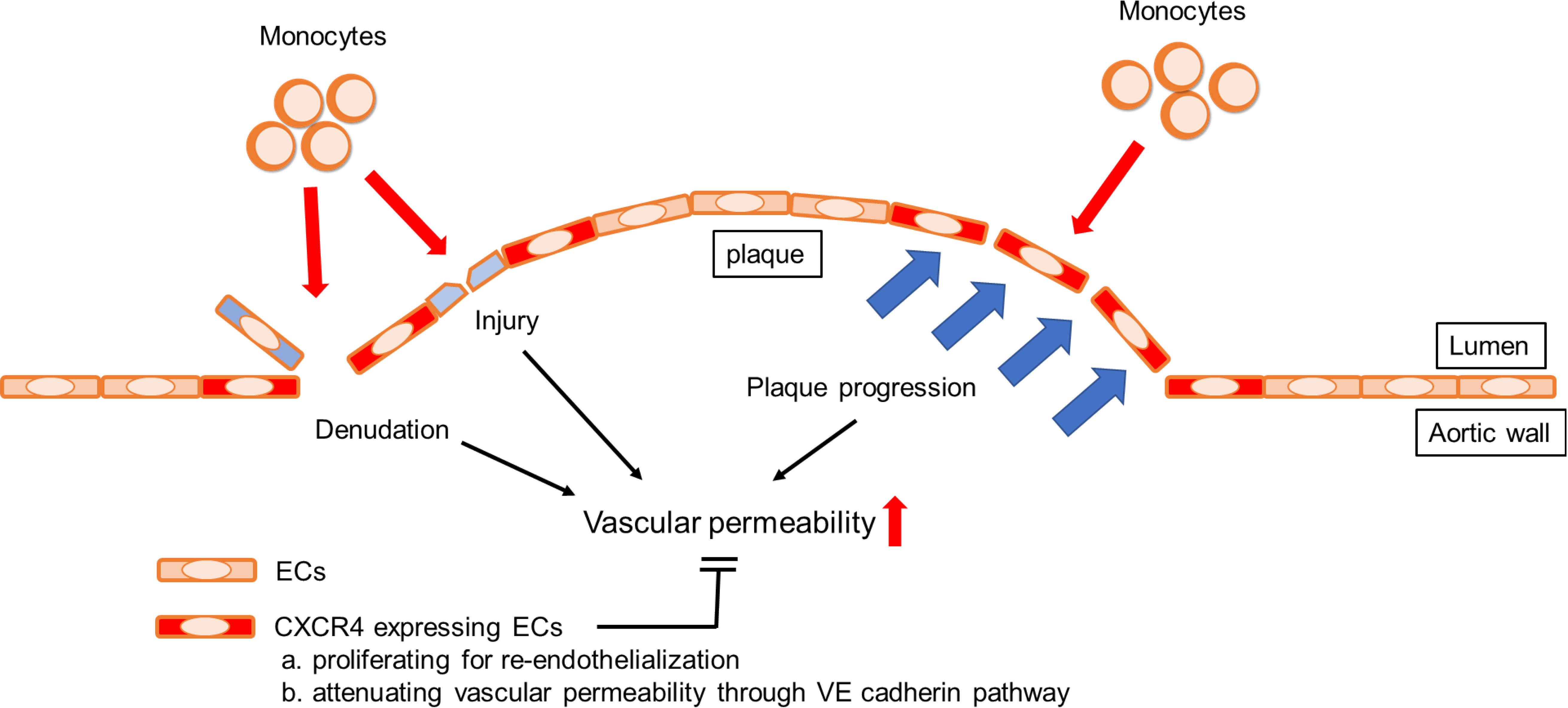
A diagram depicting factors that enhance vascular permeability and its regulation by endothelial CXCR4.

However, pathophysiological activities which enhance vascular permeability are not completely compensated by the upregulation of CXCR4 and signs of ongoing injury include monocyte recruitment. The potential utility of this tracer in a clinical setting may include detection of very early plaques even before atherosclerosis is formed, which is currently difficult, because monocyte recruitment or increased endothelial permeability is important for the initiation of atherosclerosis (29, 46, 47). Given that impairment of re-endothelialization is the important factor inducing restenosis after percutaneous coronary intervention (48, 49), PET imaging of endothelial CXCR4 targeting tracer may also be useful to predict fatal complications after coronary intervention. Even though nowadays restenosis after stenting is drastically reduced by drug-eluting stent, very late stent thrombosis is arising as a too-often fatal problem instead, and this problem is also caused by delayed re-endothelialization (49–51). Furthermore, this tracer might be useful to detect unstable plaque by recognizing endothelial denudation which is one of the features of plaque vulnerability (52, 53) although further human study is necessary because of lack of an animal model of plaque rupture. For whatever such clinical utilization, selected high risk patients would be a target of the PET scanning in terms of the cost and radiation exposure.

It is important to point out that an argument has been that a different tracer targeting CXCR4, ^68^Ga-Pentixafor, has been reported to have the capacity to penetrate plaques and reach deeper macrophages therein (41). ^68^Ga-Pentixafor is, in fact, a smaller molecule than our chemokine receptor-targeting tracers and thus may have distinct bioavailability. We would argue that a tracer with limited bioavailability may have some advantages for the purposes of examining endothelial status of plaques. Thus, a tracer with limited access to deeper area of plaques may behave differently than ^68^Ga-Pentixafor. Future studies will be needed to address this point.

The source of the ligand for CXCR4, CXCL12 is another interest for future studies. Recently, elevated blood CXCL12 level has been identified as a risk factor of coronary artery disease (54, 55) and endothelial specific CXCL12 deletion was reported to accelerate plaque progression in murine atherosclerosis model (55), which is the opposite effect to endothelial CXCR4. However, CXCL12 has another receptor CXCR7 expressed on ECs (34) which regulates systemic CXCL12 level (56). Considering these complexities, an important future direction will be to investigate the biology of CXCL12 in the endothelial injury response. It may be the case that the local availability of CXCL12 determines the pace at which the injured endothelium returns to normalcy, thereby regulating the period of time that a plaque might be most at risk. In summary, we report that ^64^Cu-DOTA-vMIP-II PET tracer identifies monocyte recruitment in part by directly binding to monocytes, but also because it identifies areas of injured endothelium through binding to CXCR4. We thus believe that this tracer has potentially ideal characteristics to identify plaques that may need clinical attention. Furthermore, because CXCR4 expressing endothelial cells are positioned in key locations, like the border of inflammatory plaques, future efforts to combine CXCR4 targeting with state-of-the-art nanotherapeutic approaches (57) may be an especially appealing way to enrich therapeutics to the most pathologic atherosclerotic lesions.

## Methods

### Mice

Mice were housed in specific pathogen-free animal facilities, maintained by Washington University School of Medicine. *Apoe^-/-^* (B6.129P2-*Apoe^tm1Unc^*/J, 002052) mice were purchased form Jackson Laboratories, then bred and maintained at Washington University School of Medicine. Diet was switched to high-fat-diet (HFD) (Teklad) containing 21% milk fat and 0.15% cholesterol from 5-6 weeks of age for 10 weeks to accelerate atherosclerotic plaque formation. All experiment procedures were performed in compliance with guidelines set forth by the NIH Office of Laboratory Animal Welfare and approved by the Institutional Animal Care and Use Committee of Washington University.

### Autoradiography of whole thoracic aortas

^64^Cu-DOTA-vMIP-II tracer was synthesized and radiolabeled as we previously reported (20). Whole thoracic aorta of *Apoe^-/-^* on HFD for 10 weeks was collected 3 h after intravenous injection of ^64^Cu-DOTA-vMIP-II tracer via tail vein and surrounding tissues were cleaned up. Collected aortas were cut, opened, washed with PBS, placed on a charged phosphor screen and exposed overnight. Images were obtained with a GE Typhoon FLA 9500 Variable Mode Laser Scanner at 50 micron resolution. After taking autoradiography images, samples were fixed with 4% paraformaldehyde (PFA) and stained with oil red O (ORO) to assess distribution of atherosclerotic plaques. Specifically, aortas were pretreated with 100% propylene glycol (Sigma-Aldrich) for 15 min, then incubated with ORO solution (Sigma-Aldrich) for 3 h at RT. They were washed with 85% propylene glycol, and then washed 3 times with PBS. After staining, aortas were placed between 2 clear sheets and imaged by M205 FA stereoscope system (Leica).

### Positron Emission Tomographic (PET) Imaging and biodistribution of PET tracer

PET imaging: Mice were anesthetized with isoflurane and injected with 3.7 MBq of ^64^Cu-DOTA-vMIP-II or ^64^Cu-DOTA-FC131 in 100 µL of saline via the tail vein. Small animal PET scan (40-60 min dynamic scan) were performed on either microPET Focus 220 (Siemens, Malvern, PA) or Inveon PET/CT system (Siemens, Malvern, PA). The microPET images were corrected for attenuation, scatter, normalization, and camera dead time and co-registered with microCT images. All of the PET scanners were cross-calibrated periodically. The microPET images were reconstructed with the maximum a posteriori (MAP) algorithm and analyzed by Inveon Research Workplace. The uptake was calculated as the percent injected dose per gram (%ID/g) of tissue in three-dimensional regions of interest (ROIs) without the correction for partial volume effect.

Post-PET biodistribution: At 3 h post injection of PET tracer, the mice were anesthetized with inhaled isoflurane prior to euthanasia by cervical dislocation. Organs of interest were collected, weighed, and counted in Beckman 8000 gamma counter (Beckman, Fullterton, CA). Standards of each tracer were prepared and counted with the biodistribution samples together to calculate the percentage of the injected dose per gram of tissue (%ID/g).

### Generation of PET tracers and dye-conjugated tracers

^64^Cu-DOTA-vMIP-II was generated as previously described (20). The FC131 peptide (cyclo[2-Nal-Gly-d-Tyr-NMe-d-Orn-Arg]) was synthesized by CPC Scientific (Sunnyvale, CA). DOTA-NHS-ester was purchased from Macrocyclics (product #: B-280). mPEG®_15_-amine was obtained from Quanta BioDesign, Ltd. (Product #: 10298). Sulfo-Cyanine5 NHS ester (sulfo-Cy5-NHS-ester) was purchased from Lumiprobe (Catalog #: 13320). CF640R Succinimidyl Ester (CF640R-NHS-ester) was obtained from Biotium (Catalog #: 92108). The ^64^Cu radionuclide (half-life = 12.7 h, β^+^ = 17%, β^-^ = 40%) was produced by the Mallinckrodt Institute of Radiology (MIR) Cyclotron Facility at Washington University School of Medicine with specific activity of 518 ± 281 GBq μmol^-1^. All other solvents and chemicals were obtained from Fisher Scientific, Sigma-Aldrich, or TCI America and were used without further purification.

*Synthesis of DOTA-FC131*. To a solution of FC131 (NMe-D-Orn^4^, 1.2 mg, 1.7 μmol) and DOTA-NHS-ester (13 mg, 17 μmol) in 200μL of dimethylformamide (DMF), N,N-diisopropylethylamine (DIPEA, 4.4 μL, 26 μmol) was added at RT. After overnight incubation at 4°C, the reaction mixture was precipitated into diethyl ether twice. The resulting white solid was collected by centrifugation, dissolved in water, purified by RP-HPLC with a H_2_O/acetonitrile [with 0.1% trifluoroacetic acid (TFA)] solvent system, and lyophilized to obtain a white solid (yield = 1.6 mg). ESI-HRMS m/z: [M + H]^+^ calcd for C_52_H_74_N_13_O_13_, 1088.5529; found, 1088.5528, [M + 2H]^2+^ calcd for C_52_H_75_N_13_O_13_, 544.7804; found, 544.7805. *Radiolabeling of DOTA-FC131 with ^64^Cu.* ^64^CuCl_2_ and DOTA-FC131 were incubated at a ratio of 1 µg precursor per 37 MBq ^64^CuCl_2_ in 50 μL of 20 mM sodium acetate buffer (pH 7) for 45 min at 45°C. The radiochemical purity (RCP) of ^64^Cu-DOTA-

FC131 was determined using radio-HPLC with a H_2_O/acetonitrile (with 0.1% TFA) solvent system. For animal studies, ^64^Cu-DOTA-FC131 with above 95% RCP was administered.

*Synthesis of DOTA-PEG*. DOTA-NHS-ester (1 mg), mPEG®_15_-amine (1.4 mg), and DIPEA (1.4 μL) in 100μL of acetonitrile were stirred overnight at room temperature. The resulting reaction mixture was purified to afford DOTA-PEG using RP-HPLC with a H_2_O/acetonitrile (with 0.1% TFA) solvent system. MALDI-TOF m/z: [M + H]^+^ calcd, 1078.6; found, 1078.4.

*Radiolabeling of DOTA-PEG with ^64^Cu*, ^64^CuCl2 (ca. 550 μCi or 20 MBq) and DOTA-PEG (0.5 nmol) were incubated in 50 μL of 0.1 M ammonium acetate buffer for 30 min at 90°C. The RCP (>95%) of ^64^Cu-DOTA-PEG was determined using radio-HPLC with a H_2_O/acetonitrile (with 0.1% TFA) solvent system.

*Synthesis of Sulfo-Cy5-vMIP-II*. Sulfo-Cy5-NHS-ester (1 mg, 1.29 μmol) and vMIP-II peptide (5.3 mg, 0.67 μmol) were incubated overnight in 4 mL of 0.1 M phosphate buffer (pH 7) at 4°C. Sulfo-Cy5-vMIP-II was isolated by RP-HPLC using a H_2_O/acetonitrile (with 0.1% TFA) solvent system. After lyophilization, 10mL of water was added to the resulting blue powder. After centrifugation (5,000 g × 5 min) to collect soluble fraction, supernatant was lyophilized to yield a blue powder. MALDI-TOF *m/z*: found, 8593 (vMIP-II conjugated with 1 sulfo-Cy5 moiety) and 9218 (vMIP-II conjugated with 2 sulfo-Cy5 moiety).

*Synthesis of CF640R-vMIP-II*. CF640R-NHS-ester (1 μmol) and vMIP-II peptide (3.98 mg, 0.5 μmol) were incubated overnight in 3 mL of 0.1 M phosphate buffer (pH 7) at 4°C. The resulting reaction mixture was purified by RP-HPLC using a H_2_O/acetonitrile (with 0.1% TFA) solvent system to yield CF640R-vMIP-II as a blue powder. MALDI-TOF *m/z*: found, 8784 (vMIP-II conjugated with 1 CF640R moiety) and 9599 (vMIP-II conjugated with 2 CF640R moiety).

*Synthesis of CF640R-FC131*. CF640R-NHS-ester (1 μmol), FC131 peptide (0.56 mg), and DIPEA (0.6 μL) were incubated overnight in 100 μL of DMF. The resulting reaction mixture was purified by RP-HPLC using a H_2_O/acetonitrile (with 0.1% TFA) solvent system to yield CF640R-FC131 as a dark green solid. MALDI-TOF *m/z*: [M+Na]^+^ calcd, ∼1442; found, 1438.7.

### Plaque regression model

Atherosclerotic plaque regression was induced in *Apoe^-/-^* on HFD for 10 weeks as previously described (12). Briefly, 1.0 x10^12^ GC of adeno-associated virus 8 vector encoding murine *Apoe* (AAV-mApoE vector, Vector Biolabs) was injected intravenously to complement *Apoe* expression. Only male mice were used in this model because AAV vector is much less effective in transducing female mice (58). Then, PET imaging of ^64^Cu-DOTA-vMIP-II tracer, biodistribution of the tracer in aortas, monocyte recruitment, macrophage accumulation and atherosclerotic plaque area were assessed at baseline, and 2 or 4 weeks after transduction. After PET imaging, aortas were taken out, weighted and counted in gamma counter to measure biodistribution. Hearts were fixed and embedded in OCT compound for cryosections (10μm) of aortic sinuses. Monocyte recruitment was assessed by counting bead-labeled monocytes in atherosclerotic plaques. In more detail, Ly6C^hi^ monocytes were labeled by i.v. injection of 1.0 µm Fluoresbrite® Polychromatic Red Microspheres (Polysciences Inc.) 2 days before sacrifice and 3 days after i.v. injection of 200 μL of clodronate-loaded liposomes (Liposoma BV). Approximately 30% of Ly6C^hi^ monocytes are labeled by this method (Fig S1D). Macrophage accumulation in atherosclerosis was assessed by staining with anti-CD68 mAb (FA-11, Bio-Rad) and MOMA-2 (Bio-Rad). Plaque size was analyzed by ORO staining to provide contrast. Plasma total cholesterol was measured by Cholesterol E kit (Wako) according to the manufacture’s instruction.

### ORO staining and Immunohistochemistry

ORO staining was done by incubating sections in 60% isopropanol containing 0.3% ORO for 10 min at room temperature. Images were taken by Eclipse E800 microscope (Nikon). For immunofluorescence staining, sections were blocked with 3% bovine serum albumin (BSA) in the presence of 1% Triton X-100 for 1 h at room temperature. Then, sections were incubated with primary antibody solution overnight at 4°C. After washing with PBS, sections were incubated with secondary antibody solution for 90 min at RT.

Nuclei were stained with DAPI or SYTOX Green (Helix NP Green, Biolegend). Immunofluorescence images were taken by TCS SPE or SP8 microscope system (Leica). Images were analyzed using Image J software. Following antibodies were purchased and used for immunostaining: CD68 (FA-11, Bio-Rad), MOMA-2 (Bio-Rad), CD45.2 (104, Biolegend), CD31 (MEC13.3, BD Pharmingen), αSMA (1A4, Sigma-Aldrich), CXCR4 (2B11, eBioscience).

Endarterectomy samples from peripheral arterial disease were collected as de-identified tissue specimens from the Vascular Surgery Biobank Repository at Washington University (IRB protocol # 201111038). The following antibodies were purchased and used for immunostaining of human sample using staining protocols similar to those used in mouse sections: CD31 (AF806, R&D systems), CXCR4 (2B11, eBioscience or UMB2, Abcam).

### Whole mount staining of aortas

After collection, aortas were fixed in 4% PFA containing 40% sucrose overnight. Then, they were blocked with 3% BSA in the presence of 1% Triton X-100 overnight at 4°C with shaking. After blocking, aortas were incubated with primary antibodies solution overnight at 4°C with shaking. Samples were washed with PBS 3 times and incubated with secondary antibodies overnight at 4°C again. After washing with PBS, aortas were cut, open and placed in an imaging chamber. Then, confocal images were taken by TCS SP8 microscope system (Leica). When tissue clearing was necessary to observe deeper sites, samples were incubated in a series of concentration of ethanol (50, 70 and 90% for 1 h, and then 100% overnight) for dehydration after incubation with secondary antibodies. Adventitia were peeled off after dehydration. Then, aortas were incubated in methyl salicylate for 30 min for tissue clearing and placed in an imaging chamber to take confocal images. Confocal images were analyzed by Imaris software (Bitplane). Following antibodies were purchased and used for whole mount staining: CD68 (FA-11, Bio-Rad), CD45.2 (104, Biolegend), CD31 (2H8, Millipore), αSMA (1A4, Sigma-Aldrich), CXCR4 (2B11, eBioscience), Ki-67 (ab15580, Abcam).

### Flow cytometry

For the analysis of peripheral leukocyte population, blood was collected by puncture of submandibular vein. Red blood cells were lysed using a commercial RBC lysis solution (BD PharmLyse, BD Biosciences). Then, samples were incubated with antibodies for 30 min at 4°C for flow cytometric analysis. For peripheral blood analysis, following antibodies were purchased and used: CD45 (30F-11, BD bioscience and Biolegend), CD11b (M1/70, Biolegend and eBioscience), Ly6G (1A8, Biolegend), CD3ε (145-2C11, eBioscience), CD45R/B220 (RA3-6B2, Biolegend), CD115 (AFS98, eBioscience), Ly6C (HK1.4, Biolegend), CXCR4 (2B11, eBioscience)

When analyzing aortic cells, whole aorta from aortic root to bifurcation was collected, finely minced and digested with digestion buffer containing 2 mg/mL of collagenase type IV (Sigma-Aldrich), 1.2 U/mL of dispase II (Sigma-Aldrich) and 5% FBS in HBSS supplemented with 0.9 mmol/L CaCl_2_ (59). Tissues were incubated at 37°C for 45 min with inversion every 15 min. After incubation, samples were triturated by pipetting 100 times and passed through a 100 μm nylon mesh (Cell strainer, BD Bioscience). After washing with HBSS supplemented with 2% FBS twice, cells were blocked with anti-CD16/CD32 antibody (Fc block, BD Pharmingen) for 20 min and then labeled with antibodies for 30 min at 4°C. Live cells were identified using SYTOX green (Helix NP Green, Biolegend) or Fixable Viability Dye eFluor 455UV (eBioscience). For Ki-67 staining, cells were fixed and permeabilized overnight using foxp3/transcription factor staining buffer (eBioscience) at 4°C. For MCP-1 intracellular staining, cells were incubated with 5.0 μg/mL of Brefeldin A (Enzo Life Science) for 5 h at 37°C, fixed by methanol-free 4% PFA for 20 min at 4°C, and then permeabilized with BD Perm/Wash buffer (BD Biosciences) for 15 min at 4°C. When analyzing aortic ECs, doublets were not excluded to include all possible cell sizes and morphologies. Following antibodies were purchased and used for aortic cell staining: CD45 (30F-11, BD Bioscience and Biolegend), CD31 (MEC13.3, Biolegend), ICAM2 (3C4, BD bioscience and eBioscience), αSMA (1A4, Sigma-Aldrich), CD11b (M1/70, eBioscience), Ly6G (1A8, Biolegend), Ly6C (HK1.4, Biolegend), CXCR4 (2B11, eBioscience), CXCR7 (11G8, R&D systems), Ki-67 (ab15580, Abcam), MCP-1 (2H5, Biolegend).

### Depletion of peripheral neutrophils and Ly6C^hi^ monocytes

One day before PET imaging study, 500 μg of anti-Ly6G mAb (clone 1A8, BioXcell) to deplete peripheral neutrophils, 250 μg of anti-Gr-1 mAb (clone RB6-8C5, BioXcell) to deplete both neutrophils and Ly6C^hi^ monocytes or PBS as control was injected into *Apoe^-/-^* mice intraperitoneally (60). Peripheral blood was collected by submandibular bleeding 1 h before PET imaging study to assess depletion efficiency by flow cytometry.

### Measurement of glycosaminoglycan (GAG) content in aortas

Aortas were collected from *Apoe^-/-^* mice on HFD 2 weeks after AAV-mApoE or PBS treatment. Then, sulfated GAG content and its subtypes were measured by Blyscan Glycosaminoglycan Assay kit (biocolor) according to the manufacture’s instruction.

### Assessment of binding of dye-conjugated vMIP-II tracer to U87 cells expressing either *CXCR4 or CXCR7*

U87 cells were obtained from NIH AIDS Reagent Program. This cellular background was chosen for its absence of endogenous CXCR4 and CXCR7, as previously demonstrated (19). pBABE.puromycin vectors (Addgene) encoding CXCR4 and CXCR7 were transfected into U87 cells and cells stably expressing the receptors were obtained following puromycin selection and subsequent single-cell sorting using BD FACSAria II cell sorter (BD Biosciences) (61). Receptor surface expression levels were verified by flow cytometry using CXCR4- and CXCR7-specific mAbs (12G5, R&D Systems and 8F11-M16, BioLegend) on a BD FACS Fortessa cytometer (BD Biosciences).

Then, U87.CXCR4 or U87.CXCR7 were distributed into 96-well plates (15 × 10^4^ cells per well) and incubated with Cy5- or CF640R-conjugated vMIP-II at concentrations ranging from 1 nM to 1 µM for 90 min on ice. Non-specific binding was evaluated on CXCR7- and CXCR4-negative U87 cells and subtracted. Dead cells were excluded using Zombie Green viability dye (BioLegend). vMIP-II binding was quantified by mean fluorescence intensity on a BD FACS Fortessa cytometer (BD Biosciences).

### Assessment of vascular permeability with Evans Blue

To assess vascular permeability in atherosclerotic aortas, 200 μL of 0.5% Evans Blue solution was injected into *Apoe^-/-^* mice intravenously. Then, 30 min later, they were euthanatized by CO_2_ inhalation and thoracic aortas were collected after perfusion with ice cold PBS. Collected aortas were placed between 2 clear sheets and imaged by M205 FA stereoscope system with Cy5 channel (Leica). CXCR4 co-staining was done after fixation using whole mount staining protocol described above.

### Statistics

Data are presented as mean ± SEM. Statistical comparisons were performed using unpaired two-tailed Student’s *t* test. When comparing 2 populations in the same mouse, paired *t* test was used. In dealing with experiments included more than two groups, statistical significance was tested by one-way analysis of variance with the Tukey’s post hoc test. The level of significance was set at a probability value of less than 0.05.

## Author contributions

OB designed and performed the experiments, analyzed the data and wrote the manuscript. AE acquired the preliminary data. DS, HL and LD assisted in performing the experiments handling radioactive tracers. GSH and XZ generated the radioactive and dye-conjugated tracers. MS and AC performed and analyzed the binding assay for CXCR4 and CXCR7. YL and GJR conceived the study and assisted in designing the experiments. YL supervised and analyzed the PET imaging experiments. GJR supervised the whole study and wrote the manuscript.

## Acknowledgments

This work was funded by R01 HL125655, R35 HL145212, and P41 EB025815 to YL; by DP1DK109668, R37AI049653, HL118206, and Washington University internal funds to GJR; and Luxembourg Institute of Health (LIH) MESR and Luxembourg National Research Fund (FNR) (PRIDE-11012546 “NextImmune”). OB was the recipient of a one-year fellowship from the Banyu Life Sciences Foundation International. His present address is University of Kyoto, Japan. AE was supported by T32 HL07081. His present address is Missouri Baptist University, Saint Louis, MO. Resources from the Diabetes Research Center were also utilized and funded by P30 DK020579. We thank Mary Wohltmann for expert assistance with the mouse colony.

